# Mapping Human Pluripotent Stem Cell-Derived Erythroid Differentiation by Single-Cell Transcriptome Analysis

**DOI:** 10.1101/859777

**Authors:** Zijuan Xin, Wei Zhang, Shangjin Gong, Junwei Zhu, Yanming Li, Zhaojun Zhang, Xiangdong Fang

## Abstract

There is an imbalance between the supply and demand of functional red blood cells (RBCs) in clinical applications. This imbalance can be addressed by regenerating RBCs using several *in vitro* methods. Induced pluripotent stem cells (iPSCs) can handle the low supply of cord blood and the ethical issues in embryonic stem cell research and provide a promising strategy to eliminate immune rejection. However, no complete single-cell level differentiation pathway exists for the iPSC-derived RBC differentiation system. In this study, we used iPSC line BC1 to establish a RBCs regeneration system. The 10× genomics single-cell transcriptome platform was used to map the cell lineage and differentiation trajectories on day 14 of the regeneration system. We observed that iPSCs differentiation was not synchronized during embryoid body (EB) culture. The cells (day 14) mainly consisted of mesodermal and various blood cells, similar to the yolk sac hematopoiesis. We identified six cell classifications and characterized the regulatory transcription factors (TFs) networks and cell-cell contacts underlying the system. iPSCs undergo two transformations during the differentiation trajectory, accompanied by the dynamic expression of cell adhesion molecules and estrogen-responsive genes. We identified different stages of erythroid cells such as burst-forming unit erythroid (BFU-E) and orthochromatic erythroblasts (ortho-E) and found that the regulation of TFs (e.g., TFDP1 and FOXO3) is erythroid-stage specific. Immune erythroid cells were identified in our system. This study provides systematic theoretical guidance for optimizing the iPSCs-derived RBCs differentiation system, and this system is a useful model for simulating *in vivo* hematopoietic development and differentiation.

## Introduction

The substantial gap between the supply and demand for blood has always been a serious clinical practice problem [1]. Artificial blood is an essential means of solving this worldwide shortage, and obtaining functional red blood cells (RBCs) is the key to artificial blood generation [2]. RBC regeneration *in vitro* has been an important research direction in the blood research field for many years [3, 4]. In the current study, the primitive materials that can be used for RBC regeneration are mainly cord blood hematopoietic stem cells (HSCs) [5], embryonic stem cells (ESCs) [6, 7], and induced pluripotent stem cells (iPSCs) [6, 8]. The regeneration technology of cord blood HSCs is relatively mature; however, some problems come with this technology, such as difficulty obtaining materials, significant differences among individuals, the high price of this technology, and ESCs are subject to ethical restrictions [9]. Human iPSCs can be differentiated into various cell types *in vitro*, providing a model for basic research and a source of clinically relevant cells [10–12]. Although the derivation of iPSC lines for patients has now become a routine technique, how to accurately differentiate iPSCs into the desired cell types is still a major challenge [13, 14].

Decades of research have shown that *in vitro* hematopoietic differentiation is closely related to *in vivo* development [15]. Hematopoiesis originates from the *FLK1*+ (*KDR*) lateral mesoderm, and some of the specialized cells can be induced to form vascular hemangioblasts by the transcription factor (TF) ETV2 [16, 17]. These cells can differentiate into the blood and vascular precursor cells, during which ETV2 is regulated by the BMP and WNT signaling pathways [18, 19]. At this time, the primitive RBCs that express embryonic globin (ε-globin; *HBE*) are produced, which is the first wave of hematopoiesis in the yolk sac [20]. Hemangioblasts differentiate into endothelial cells and specialize in hematopoietic endothelial cells that produce erythroid-myeloid progenitors (EMPs). EMPs can differentiate into most of the myeloid and into definitive RBCs that can simultaneously express embryonic (ε-globin; *HBE*), fetal (γ-globin, *HBG1* and *HBG2*), and adult (β-globin, *HBB*) globin; this differentiation represents the second wave of hematopoiesis in the yolk sac [21]. HSCs can also be directly produced through the endothelial-hematopoietic transition (EHT) induced by GATA2 and RUNX1 [22–24]. This occurs in the aorta-gonad-mesonephros (AGM) region that maintains lifelong hematopoietic differentiation and the hematopoietic cycle [25], and these cells can differentiate into definitive RBCs that express only adult globin and are regulated by signaling pathways such as VEGF and HIF [26, 27]. Hematopoietic differentiation is the differentiation process of HSCs to form various types of blood cells [28]. After the differentiation and activation of HSCs, it is necessary to go through the cell fate determination program regulated by TFs GATA1, KLF1, and other TFs to enter the erythroid differentiation process [29]. The generation of RBCs is an elaborate multistep process involving the differentiation of early erythroid progenitor cells into definitive RBCs, which require spatiotemporal specific completion of globin synthesis and assembly [30, 31], iron metabolism [32, 33], heme synthesis [34], cell denucleation, and other processes, eventually forming intact functional RBCs [35]. However, the molecular and cellular mechanisms involved in these processes are still poorly understood.

Based on the existing molecular mechanisms of hematopoietic development and erythroid differentiation, scientists simulate embryonic hematopoiesis [36] using the spin EB method to regenerate hematopoietic stem progenitor cells (HSPCs) *in vitro* and then induced the differentiation of RBCs from iPSCs. However, the differentiation efficiency and denucleation rate are low, and adult globin expression is limited; these drawbacks limit the clinical availability [37]. Simultaneously, iPSC differentiation is an elaborate process, and flow cytometry and immunostaining have been used to determine the cell types during iPSC differentiation culture. However, these methods are limited by the number of fluorescent probes used, and it is not possible to solve important problems, such as the cell composition and differentiation path of the iPSC differentiation system under high-resolution conditions. Recently, single-cell transcriptome technology has been able to capture and identify rare and transient cell types, determine the spatial or temporal localization of cells, and reconstitute gene regulatory networks, helping scientists understand the mechanisms by which development and cell fate are determined [38]. In recent years, there have been many single-cell hematopoiesis studies *in vitro* [39, 40]. By single-cell sequencing of embryoid bodies, researchers found that naïve H9 ESCs have a stronger hematopoietic capacity than primer H9 ESCs [41] and illuminated the heterogeneity of pluripotent stem cell-derived endothelial cell differentiation [42]. Single-cell sequencing of day 29 erythrocytes belonging to the iPSC-derived erythroid differentiation system revealed that cells expressing β-globin showed reduced transcripts encoding ribosomal proteins and increased expression of ubiquitin-proteasome system members [43]. The lack of a complete single-cell transcriptome map, as iPSCs differentiate into RBCs, does not allow scientists to guide iPSCs to produce functional RBCs. Therefore, in this study, we established an iPSC-derived erythroid differentiation system and obtained a dynamic transcriptional map of cell differentiation through high-resolution single-cell transcriptomics sequencing. The cell map of the *in vitro* iPSC-derived RBC differentiation system and the *in vitro* differentiation trajectory of the intact iPSCs to RBCs were mapped for the first time. iPSCs undergo two transformations, during the first transformation, EHT, withdraw from pluripotency into lateral plate mesoderm and differentiate into hematopoietic endothelial cells and then into HSPCs, accompanied by cell adhesion molecules expression fluctuation. During the second transformation, HSPCs differentiate into erythroid progenitor cells and erythroid differentiation, estradiol promotes erythroid progenitor cell proliferation and inhibits erythroid differentiation. This study has an important guiding significance for analyzing the basic fate of cells and the gene network structure, as well as for designing more effective hematopoietic and erythropoiesis strategies for regenerative medicine.

## Results

### iPSC cell line-derived HSPC production and erythropoiesis

Depending on the spin EB culture method, we used the iPSC cell line BC1 to collect the suspension cells (SCs) through the cell strainer in the system after 14 d of EB culture [8, 44, 45] (Figure 1A). The established culture system can effectively produce SCs. Cell count analysis showed that the number of SCs on day 14 was approximately 10-fold higher than the number of iPSCs on day 0 (Figure 1B). Flow cytometry analysis showed that approximately 25.9% ± 3.05% of the SCs expressed cell surface marker CD34, approximately 27.08% ± 1.63% of the SCs expressed cell surface marker CD45, and approximately 22.9% of the SCs expressed both CD34 and CD45, indicating that we collected a certain percentage of HSPCs from the SCs (Figure 1C). To detect hematopoietic differentiation’s potency, we performed colony-forming unit (CFU) assays on the collected SCs from day 14. After two weeks of culture, differential blood lineage colonies were observed and counted under the microscope (Figure 1D and E). It suggested that the SCs have the potential to differentiate into various blood lineage cells. Subsequently, we attempted to induce HSPCs in SCs toward erythroid differentiation using serum-free medium (SFM) [8]. By assessing changes in expression of the cell surface markers CD71 and CD235a at different time points (Figure 1F), we found that on day 14 of the EB stage, the proportion of CD71 and CD235a double-positive cells in SCs exceeded 30%, indicating that a portion of cells in the system entered the erythroid differentiation stage. On day 28, more than 95% of cells expressed both erythrocyte markers, CD71 and CD235a (Figure 1F). qPCR assay showed that the expression of *HBB* and *HBG* increased significantly during the entire system differentiation (Figure 1G). Wright-Giemsa staining analysis showed that most cells were at the terminal erythroid differentiation stage and some cells were denucleated RBCs (Figure 1H). In summary, we successfully constructed an integrated *ex vivo* hiPSCs derived erythroid differentiation system by spin EB technology using the iPSC cell line BC1, as well as obtained and induced HSPCs to enter the erythroid differentiation process, eventually producing denucleated RBCs. Though we have constructed an erythroid cell differentiation system, we still do not understand what kind of molecular changes of the system’s cells have undergone during the series of processes from pluripotent stem cells to erythroid cells, and what factors regulate the process of molecular changes experienced by cells. Therefore, to clarify the above problem, we collected cells that contain various cell types on day 14 at the middle time point of the differentiation system and performed single-cell transcriptome sequencing.

**Figure 1.**
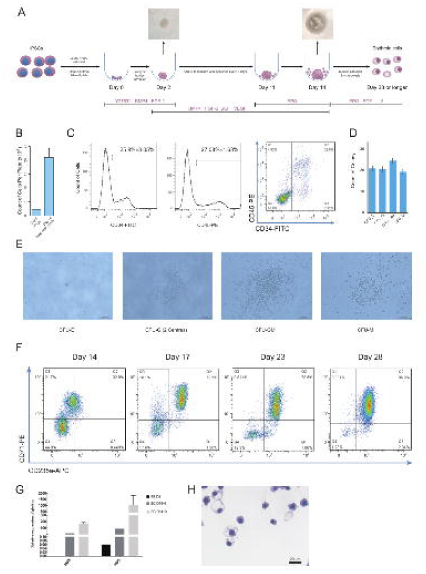
Identification of iPSC-derived HSPCs and erythroid cells. **A**. Schematic diagram of the protocol used to generate HSPCs and erythroid cells from iPSCs *in vitro*. **B**. Histogram of iPSC counts on day 0 and collected SCs on day 14 per round-bottom 96-well plate (error bar = mean ± SD). **C**. Percentage of CD34^+^ and CD45^+^ cells in the total SCs (mean ± SD). **D**. Count of colonies that grew from SCs collected on day 14. **E**. The morphology of hematopoietic colony formation units grown from SCs. Scale bar = 1000 μm. **F**. Flow cytometry results showing cells expression of CD71 and CD235a on day 14, 17, 23, and 28 during the erythroid differentiation of SCs. **G**. qPCR of β-chain and γ-chain globin expression on day 4, day 14 + 3, and day 14 + 9 during the entire system differentiation. **H**. The morphology of erythroid cells on day 28. Scale bar = 20 μm.

### Cell composition of the iPSC-derived RBC differentiation system

To construct a scRNA-seq library, we isolated and selected EB cells (EBCs) and SCs on day 14, and constructed a scRNA-seq library based on 10× Genomics platform. After rigorous quality control analysis of the data by Seurat and the removal of double cells droplets using R package DoubletFinder [46, 47], we obtained 3660 cells from batch 1 and 6538 cells from batch 2, given 10,198 high cells for downstream analysis (Figure S1A). On average, we detected 3523 informative genes and 17,825 unique molecular identifiers in each cell (Table S1). The correlation analysis of the expression pattern indicated that our 2 batch samples were positively correlated (Spearman coefficient = 0.904, *P* < 0.001) (Figure S1B). We then clustered cells based on Seurat’s K-nearest neighbor method, annotated cells based on R package SingleR [48] and published datasets [49, 50], and visualized cells based on uniform manifold approximation and projection (UMAP) [51] (Figure 2A and B; Figure S2 and S3).

**Figure 2.**
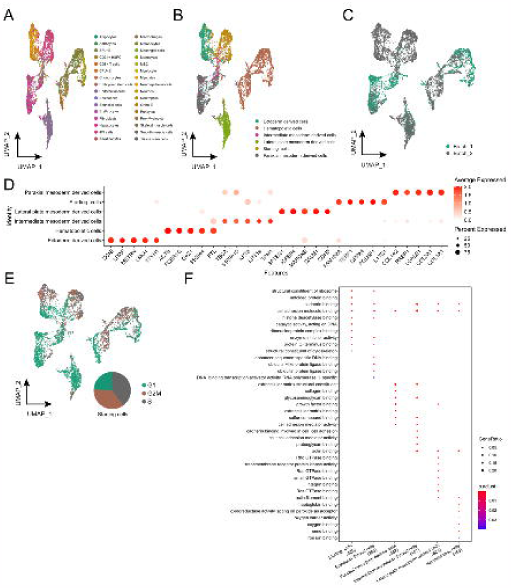
Cell composition in the iPSCs-derived system. **A**. UMAP plot of cells labeled by batches. B. UMAP visualization of EBCs and SCs. Cells were labeled by SingleR annotation results. C. UMAP plot of cells labeled by cell classifications based on starting cells or derived from different germ layers. **D**. The dot plot of signature genes for each cell classification. **E**. UMAP visualization of cell cycle analyzed by Seurat. The pie chart showed the percentage of cells in different cell cycles in the starting cells. Phases of cells indicated by different colors, G1 in green, G2M in brown, and S in gray. **F**. GO term enrichment results of signature genes from each cell classification.

We classified cells into 6 categories from the cells belonging to starting cells or coming from different germ layers. iPS cells and ESCs were defined as starting cells (n = 1705); astrocytes, neuroepithelial cells, and neurons were classified as ectoderm-derived cells [52, 53] (n = 1529); epithelial and mesangial cells were classified as intermediate mesoderm-derived cells [54, 55] (n = 413); adipocytes, fibroblasts, myocytes, mesenchymal stem, skeletal muscle, smooth muscle, and tissue stem cells were classified as paraxial mesoderm-derived cells [56, 57] (n = 1934); endothelial cells and pericytes were classified as lateral plate mesoderm-derived cells [58] (n = 1561); CD34^+^ HSPC, pro-myelocyte, myelocyte, macrophages, monocytes, neutrophils, CD8^+^ T cells, burst-forming unit erythroid (BFU-E), colony-forming unit erythroid (CFU-E), erythrocytes, and orthochromatic erythroblasts (ortho-E) were classified as hematopoietic cells (n = 3056) (Figure 2C; Figure S4A).

To prove the cell classifications’ results reliability, we identified the signature genes of each cell classification and calculated the Spearman coefficient between different cell classifications (Figure 2D; Figure S4B). It appears that the expression pattern of each cell classification was specific. We also performed cell cycle analysis based on the expression pattern of genes using Seurat and visualized the cell cycle analysis results into the UMAP dimension reduced map. We noted that among 10,198 cells, 2390 cells were in the S phase, 2076 cells were in the G2M phase, 5732 cells were in the G1 phase (Figure 2E). Specifically, we found that the cell cycle pattern was highly heterogeneous in different cell classifications, especially more than three-quarters of the starting cells in the G2M/S phase (Pie chart in Figure 2E). Based on the signature genes of 6 cell classifications, we performed the Gene Ontology (GO) enrichment analysis using R package clusterProfiler to gain insight into the biological processes involved in the EB formation and cell differentiation process [59]. It is impressive that cell adhesion-related pathways were present in all cell classifications (Figure 2F).

### ESAM affects hematopoiesis through cell-cell interactions

We identified that cell adhesion-related pathways may play an essential regulatory role during EB formation to hematopoiesis, and next analyzed the mode of cell adhesion between cells in our system. We used CellChat[60] to perform cell-cell contact analysis and evaluate the regulatory function of cell adhesion-related pathways in our system. We found that the contact score between the lateral plate mesoderm-derived cells, and other cells were the highest (Figure 3A), and the most significant cell-cell contact signal corresponding to the lateral plate mesoderm-derived cells was the contact between endothelial-specific adhesion molecule (ESAM) ligand and ESAM receptor (Figure 3B; Figure S5A and B). ESAM pathway is an autocrine signaling pathway, and ESAM was expressed and secreted by lateral plate mesoderm-derived cells, and then acted on itself and played a particular regulatory role in hematopoietic cells (Figure 3C and D). The lateral plate mesoderm-derived cells were mainly composed of endothelial cells (Figure 3D). Previous studies have shown that expression of the ESAM enhances the tight junctions between the arterial endothelium, which is not conducive for producing HSCs [61]. We performed bulk RNA-seq on SCs collected at day14 and the iPSC line BC1, and compared them with published HSPC transcriptome data from human cord blood [62]. We found that SCs are significantly different from iPSCs, and the hematopoietic lineage-related gene expression patterns are similar to HSPCs and exhibit HSPC-like characteristics (Figure S5C and D). Compared with BC1, SCs also down-regulated the expression of cell adhesion-related genes (Figure S5E), which may be the direct cause of SCs separation from EBs, and necessary demand for hematopoiesis.

**Figure 3.**
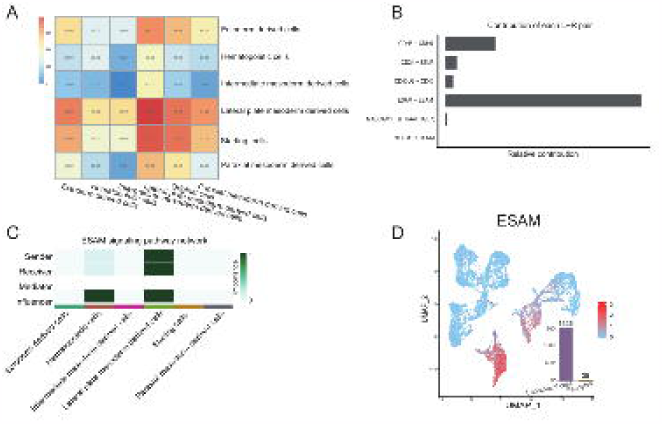
Cell-cell contacts analysis. **A**. Heatmap of cell-cell contact scores analyzed by CellChat. **B**. Contribution of each L-R pair in CDH5, CD34, CD40, ESAM, MADCAM, and PTPRM signaling pathway. **C**. The role of each classification cell in ESAM signaling pathway. **D**. UMAP plots of every single-cell *ESAM* expression level and cell counts bar plot of endothelial cells and pericytes in lateral plate mesoderm-derived cells.

### Regulons networks in the iPSC-derived RBC differentiation system

To clarify regulons (i.e., TFs and their target genes) networks from EB formation to hematopoiesis in our system, we used the TFs included in the detected data to reduce dimension and mapped the SingleR cell annotation results to the new TFs dimension atlas. The R package, SCENIC, was used to calculate the area under the curve (AUC) scores of regulons [63]. The mapping data showed that the cell differentiation trajectory revealed by TFs is consistent with the clustering by signature genes (Figure 4A), and the same cell classification still clustered together. According to the AUC scores, regulons’ activity has significant cell classification specificity (Figure 4B). Based on regulons’ activity pattern, we constructed the interaction network of cell classification-specific regulons and their target genes based on Cytoscape. We observed that part of the target genes of TFs that were activated in the same cell classification was overlapped, suggesting that the regulation of TFs was synergistic in our system (Figure 4C). We performed the GO enrichment analysis for target genes of TFs by the clusterProfiler (Figure S6) and mapped the activation position of regulons in the UMAP atlas (Figure S7). Specifically, we observed that MECOM, ETS1, ELK3, ERG, and SOX18 showed high activities in lateral mesoderm-derived cell (Figure S6I, J, and S7), while GATA1, TAL1, RUNX1, CEBPD, MYB, and ELF1 were activated in hematopoietic cells (Figure S6K, L, and S7). CEBPD and MYB were mainly activated in myeloid cells and targeted to myeloid cell differentiation and cytokine production-related biological processes (Figure S6M, N, and S7). GATA1 and TAL1 were activated explicitly in erythroid cells, and participate in erythrocyte differentiation, homeostasis, Ras signaling transduction, and GTPase activity by regulating their target genes. At the same time, RUNX1 and ELF1 exhibited activities in both myeloid and erythroid cells and enriched in cell activation and leukocyte proliferation through their target genes.

**Figure 4.**
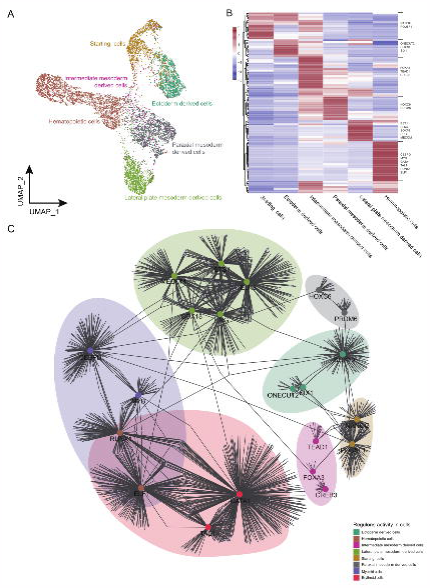
Regulons regulation analysis. **A**. UMAP map of all cells based on human transcription factors, with colors indicating six cell classifications. **B**. Heatmap plot of regulons AUC patterns related to cell classifications. The high AUC score regulons in each cell classification were listed on the right side of the plot. **C**. The network of the high AUC score regulons in each cell classification.

To identify the TFs specific to the differentiation stage during hematopoietic differentiation, we calculated the regulon specificity score (RSS) for each cell type included in hematopoietic cells separately. We noticed that erythroid cells at different stages have a unique RSS pattern, suggesting that the regulation of TFs was erythroid-stage specific (Figure S8). Especially in the erythrocytes cluster, we observed TFDP1 with the highest RSS that can heterodimerize with E2F proteins complex to control the transcriptional activity of numerous genes involved in cell cycle progression from G1 to S phase [64]. Wang et al. showed that the *kyo* mutant of TFDP1 could cause changes in the morphology and number of erythroid cells during the development of fish embryos, indicating that TFDP1 can be linked to the development of erythroid cells [65]. However, the regulatory role of TFDP1 in the development of mammalian erythroid cells has not been clarified. We speculate that TFDP1 may be related to cell number restriction during the maturation of erythroid cells.

### Cell differentiation trajectory of the iPSC-derived RBC differentiation system

To further determine the cell differentiation trajectory during the differentiation of the entire system, we used Monocle2 to order single cells and construct an entire lineage differentiation trajectory with a tree-like structure [66] (Figure 5A). The results showed that cell differentiation originated from starting cells. After differentiation into different germ layers, two branches appeared. The hematopoiesis branch was mostly hematopoietic cells, and the non-hematopoiesis branch consisted of ectoderm, intermediate mesoderm, and other cells (Figure 5B; Figure S9A). We use a pseudo-time branch heatmap to show branch-specific expression patterns along two developmental pathways (hematopoiesis *vs*. non-hematopoiesis). Specifically, we identified 264 signature genes (cluster 1) for pre-branch, including 39 TF coding genes, such as *POU5F1* and *SOX2*, enriched by glycolysis/gluconeogenesis. Subsequently, pre-branch cells were driven by RUNX1, GATA2, and 22 other TFs to enter the hematopoiesis branch (cell fate 1), and 300 signature genes (cluster 2) were identified for hematopoiesis that were enriched in hematopoietic cell lineage pathway. Oppositely, pre-branch cells were driven by SOX7, SOX17, and 41 other TFs to enter non-hematopoiesis branch (cell fate 2), and 409 signature genes (cluster 2) were identified for non-hematopoiesis that were enriched in vascular smooth muscle contraction pathway (Figure 5C and D; Figure S9B). Several research teams have proved that RUNX1 and GATA2 mark the beginning of hematopoiesis [23, 67]. SOX7 activates CDH5 promoter and hinders the binding of RUNX1 to DNA, and SOX17 mediates the repression of RUNX1 and GATA2, thereby hindering the transition to hematopoietic fate and maintaining the endothelial-arterial identity [67–69].

**Figure 5.**
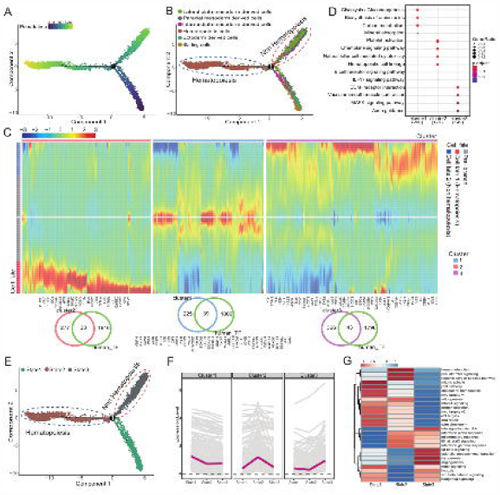
Cell pseudo-time trajectory of the iPSCs-derived RBCs regeneration system. **A**. Developmental pseudo-time of iPSCs-derived erythrocytes regeneration system by Monocle2. **B**. The arrangement of different classification cells on pseudo-time trajectory. **C**. Heatmap showing the specific gene expression patterns in hematopoietic and non-hematopoietic branches. **D**. Functional annotation analysis of each cluster gene from **C**. **E**. The arrangement of different state cells on pseudo-time trajectory. **F**. The expression of trend changes between states of each cluster gene from **D**. **G**. GSVA analysis among different cell states.

Meanwhile, Monocle2 divides cells into 3 states, pre-branch, hematopoiesis, and non-hematopoiesis branches corresponding to state 1, 2, and 3 (Figure 5E). Analysis of the expression trend of each branch signature genes in different state cells showed that each pseudo-time branch signature genes are highly expressed in its corresponding state cells (Figure 5F). The functional annotation analysis showed that state 1 cells (pre-branch) were enriched in the mitotic spindle, G2M checkpoint, mitotic prometaphase, indicating that it was in an active cell division state, and was regulated by P53, MYC, and mTOR pathways. State 2 cells (hematopoiesis branch) were enriched in heme metabolism, neutrophil degranulation, and other blood cell-related pathways, and are regulated by PI3K and IL6. State 3 cells (non-hematopoiesis branch) were enriched in angiogenesis, myogenesis, extracellular matrix organization, other mesodermal and mesenchymal cell-related pathways, and regulated by signal pathways, such as TGFβ and hypoxia (Figure 5G; Figure S9C).

To further understand the molecular characteristics of cells that entered the hematopoiesis branch, we analyzed the difference between state 2 initial cells (the first 100 cells that entered the hematopoiesis branch) and state 3 initial cells (the first 100 cells that entered the non-hematopoiesis branch) and found that state 2 initial cells had a low expression of *COL5A1*, *COL6A3*, and other collagen genes, and a high expression of *AIF1*, *SPI1*, and other genes related to myeloid differentiation (Figure S9D). The highly expressed genes retrieved from the TRRUST database are regulated by 20 TFs such as SPI1 and GATA1 (Figure S9E). At the same time, these TFs and their target genes are significantly enriched in embryonic hemopoiesis, erythroid differentiation, and myeloid cell differentiation, suggesting that our system is similar to early embryos hematopoiesis and tend to produce erythroid and myeloid cells, and were regulated by Notch signaling pathway and estrogen (Figure S9F).

### Transition from starting cell to hematopoietic progenitor cell

Previous studies have shown that HSPCs are differentiated from endothelial progenitor cells in the lateral mesoderm during embryonic development [17]. To determine how the transition from starting cells to hematopoietic cells, we conducted the pseudo-time analysis of ESCs, iPS cells, endothelial cells, and CD34^+^ HSPCs. Most of the iPS cells and ESCs fell into the pre-branch and undergo bifurcation. The cell fate 1 branch was mostly CD34^+^ HSPCs, and the cell fate two branch was mostly endothelial cells (Figure 6A and B; Figure S10A). Differential gene expression analysis was performed between the 2 branches to determine the characteristic genes in each fate cell. Interestingly, we identified hematopoietic endothelium (HE) gene set (cluster 5), including 10 TF coding genes expressed on both endothelial cells and HSPCs branch. The TF coding genes *LMO2* and *FLI1* in this gene set are the marker genes of HE during embryonic development [70, 71] . FOS, JUNB, and DAB2 are the downstream TFs of TGFβ, and TGFβ drives HE programming [72, 73] . FOS and MAP4K2 are the effect factors in the MAPK cascade that VEGF can directly regulate to establish arterial specifications of endothelial progenitor cells [74]. The gene set (cluster 5) was observed in cell adhesion molecules and tight junctions (Figure 6C).

**Figure 6.**
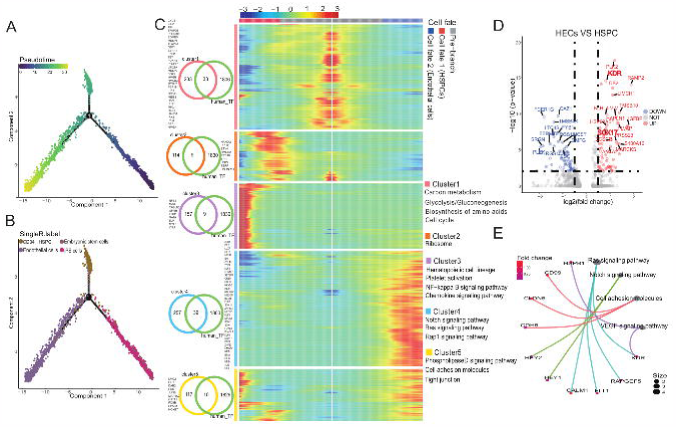
Key factors driving hematopoiesis. **A**. Developmental pseudo-time of iPS cells, Embryonic stem cells, Endothelial cells, and CD34^+^ HSPC by Monocle2. **B**. The arrangement of different type cells on pseudo-time trajectory. **C**. Heat map showing the specific gene expression patterns in endothelial cells and HSPCs branches. **D**. Volcanic plot of DEGs from gene differential expression analysis between HECs and HSPCs (*P* < 0.05, |log2 FC| < 0.5). **E**. KEGG analysis of HECs high expression gene.

HSPCs branch corresponds to state 2 cells, endothelial cells branch corresponds to state 3 cells, and HE gene set was highly expressed in state 2 and state 3 cells (Figure S10B and C). At the same time, we found that some CD34^+^ HSPCs fell in state 3, which was defined as HECs. By comparing with CD34^+^ HSPCs on state 2, we found that HECs highly expressed *KDR*, *SOX17*, and other endothelial characteristic genes and highly expressed genes were enriched in cell adhesion molecules and regulated by RAS, NOTCH, and VEGF (Figure 6D and E). We verified the expression of some cell adhesion molecules by qPCR assay combined with previous function annotation results for cell classifications marker genes and cell-cell contact pattern analysis. Results indicated that the expression of these molecules fluctuated during RBC regeneration; their expression is up-regulated during the HE formation [75] (EB day 4), while their expression is down-regulated in hematopoietic cells (SCs day 14) (Figure S10D).

### Transition from HSPC to the erythroid cell

To clarify the transition from HSPCs to the erythroid cells, we conducted a pseudo-time analysis of hematopoietic cells. Hematopoietic cells are mostly composed of CD34^+^ HSPCs, erythroid cells, monocyte-macrophages, and a small amount of lymphoid CD8^+^ T cells (Figure S11A and B). The differentiation trajectory shows fewer cells on the pre-branch, and most of the cells are ordered to the erythropoiesis and non-erythropoiesis branches (Figure 7A and B). Specifically, some BFU-E, CFU-E cells, and CD34^+^ HSPCs are distributed in the pre-branch corresponding to state 4. Non-erythroid cells (eosinophils, macrophages, monocytes, myelocytes, neutrophils, and pro-myelocytes) are mostly distributed in the non-erythropoiesis branch (cell fate 1) corresponding to state 3, and erythroid cells (BFU-E, CFU-E, erythrocytes, ortho-E) are mostly distributed in the erythropoiesis branch (cell fate 2) corresponding to state 1, 2, and 5 (Figure 7C and Figure S11C). Simultaneously, state 4 cells highly express MYB, CD34, CD45 (PTPRC), and CD44, which are yolk sac-derived myeloid-biased progenitors (YSMPs) markers, YSMPs are human yolk sac second wave hematopoietic progenitor cells. And state 4 cells almost no expression of *HOX6*, which is HSPC specific marker in the definitive hematopoietic AGM region [76], and EMP (mouse yolk sac second wave hematopoietic progenitor) marker *KIT* [77] (Figure 7D), suggesting that our system is similar to the second wave of hematopoiesis in the human yolk sac.

**Figure 7.**
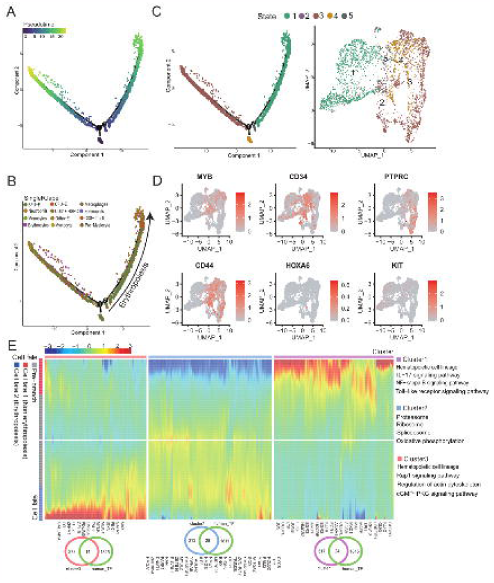
Differentiation trajectory of hematopoietic cells. **A**. Developmental pseudo-time of hematopoietic cells by Monocle2. **B**. The arrangement of different type cells on hematopoietic pseudo-time trajectory. **C**. The arrangement of different state cells on pseudo-time trajectory and UMAP plot of hematopoietic cell states distribution. **D**. UMAP plots of every single cell showing the expression level of *MYB*, *CD34*, *PTPRC*, *CD44*, *HOXA6*, and *KIT*. **E**. Heat map showing the specific gene expression patterns in erythropoiesis and non-erythropoiesis branches.

To further clarify the molecular regulation mode in the differentiation process of YSMP in the system, we performed a difference map between the 2 branches (erythropoiesis and non-erythropoiesis), YSMPs (pre-branch) were driven by TAL1, GATA1, and 12 other TFs enter the erythropoiesis branch (cell fate 2), oppositely, YSMPs were driven by CEBPD, MAF and other 22 TFs enter to non-erythropoiesis branch (cell fate 1). YSMPs (Pre-branch) identified 341 signature genes (cluster 2), enriched by oxidative phosphorylation. Interestingly, the gene set has a strong tendency for expression in erythropoiesis branch, which may be EPO driven in the system (Figure 7E and Figure S11D and E). Besides, we found that some of the erythroid cells will fall into the non-erythropoiesis branch, indicating other types of erythroid cell production in our system (Figure S11C).

### Differentiation trajectory of erythroid cells

To accurately understand the molecular activities during erythroid differentiation, we took BFU-E, CFU-E, erythrocytes, ortho-E cells to perform a pseudo-time analysis (Figure 8A). The pseudo-time result was consistent with the SingleR cell annotation result: the erythroid progenitor cells were mostly distributed at the beginning of the differentiation trajectory, and the differentiated terminal cells were mainly at the end of the trajectory (Figure 8B; Figure S12A). Furthermore, cluster analysis of the top 1000 genes with the pseudo-time scoring significance was performed to obtain the highly expressed gene set cluster 1 of erythroid progenitor cells, including the GATA1 and 36 other TFs, which were enriched in the estrogen signaling pathway. The expression trend of genes enriched in the process of erythroid differentiation is mostly initially high and then decreased. The gene set cluster 2, which was highly expressed in the erythroid cells, includes 34 TFs, such as BCL11A, which enriched cholesterol metabolism, glycosaminoglycan binding, and oxygen carrier activity (Figure 8C; Figure S12B and C). The pseudo-time analysis divides cells into 3 states; the progenitor cells correspond to state 1 in which the cluster 1 gene set was highly expressed, the erythroid cells in state 2 and three in which the cluster 2 gene set was highly expressed (Figure 8D; Figure S12D). Specifically, state 1 (erythroid progenitor cells) responds more strongly to estrogen (Figure 8E). We tested the role of estrogen by adding estradiol during the induction and differentiation process of SC cells and observed that expression of *KLF1* slightly increased in the erythroid progenitor cells, but the expression of *ALAS2*, *GATA1*, and *KLF1* in the terminal erythroid differentiation was significantly inhibited (Figure S12E). Altogether, it suggested that estrogen promotes the proliferation of erythroid progenitor cells, but may inhibit erythroid differentiation (Figure S12E).

**Figure 8.**
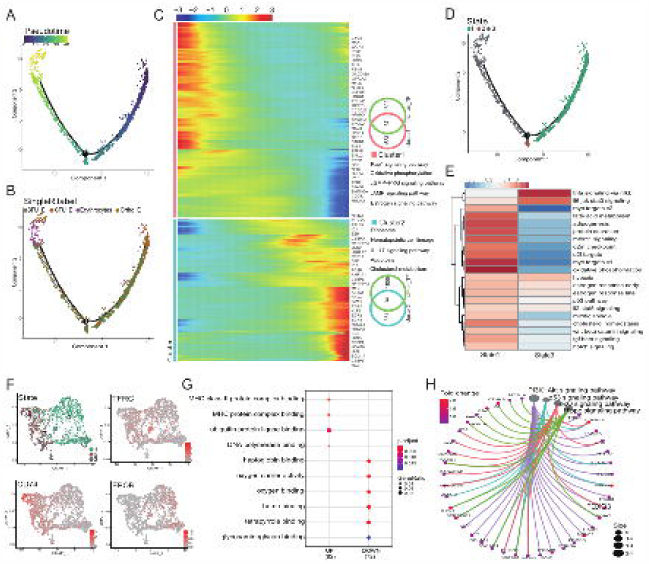
Pseudo-time trajectory of erythroid cells. **A**. Developmental process of erythroid cells by Monocle2. **B**. The arrangement of different type cells on erythroid cell pseudo-time trajectory. **C**. Heatmap showing the gene expression dynamics during erythroid differentiation. **D**. The arrangement of different state cells on erythroid cell pseudo-time trajectory. **E**. GSVA analysis between state one and three cells. **F**. UMAP plots of erythroid cell state distribution and *TFRC*, *CD74*, and *EPOR* expression level. **G**. GO analysis of DEGs between state one and three cells. **H**. KEGG Network constructed with high expression genes in ortho-E cells

Interestingly, we found that state 2 cells highly express *CD74* and lowly express *EPOR*. Compared with state 3, state 2 lowly expresses genes (e.g., *HBD* and *HBE1*) that related to oxygen carrier activity, oxygen binding and highly expresses genes (e.g., *CD81*, *C1QBP*, and other myeloid characteristic genes) that related to MHC protein complex binding genes, which can be defined as immune erythroid cells (Figure 8F and G; Figure S12F). Also, we performed a difference analysis between ortho-E and other erythroid cells (Figure S12G). Ortho-E cells had low expression of genes that are related to the ribosome (e.g., *RPS16* and *RPS13*), oxidative phosphorylation, and cell cycle. In contrast, having high expression genes that associated with PI3K-Akt and FoxO signaling pathway, among which *FOXO3* plays a crucial role in the enucleation of erythroid cells [78] (Figure 6H; Figure S12G and H).

## Discussion

With the rapid development of single-cell technology, single-cell transcriptome studies on the development of the hematopoietic lineage have been extensively reported, and the known hematopoietic theory has been revised and supplemented, but most studies focused on bone marrow, peripheral blood, cord blood, and other systems [79, 80] . Here, we used the EBs culture method verified by many laboratories to construct the RBC regeneration system derived from iPSCs [8, 44, 45, 81]. The erythroid differentiation efficiency of iPSCs generated from different type of tissues might be different, however, these iPSCs cell lines can differentiate towards erythroid cells under appropriate inducing conditions [82]. After multiple induction and differentiation experiments under the same conditions, it is clear that HSPCs in the SCs produced by the EBs can stably enter the erythroid differentiation process (data not shown). After clarifying the repeatability and stability of the system, we performed a single-cell transcriptomics study on the RBC regeneration system derived from iPSCs, in which over 10, 000 high-quality cells of the regeneration system from two batches were analyzed. The two batches of cells are positively correlated, and the mesoderm and hematopoietic cells are dominantly present in each batch. To avoid the in-between cell types in the regeneration system, we applied four datasets that are integrated by the R package SingleR, including Human Primary Cell Atlas reference dataset, BLUEPRINT and Encode reference dataset, Database of Immune Cell Expression reference dataset and Novershtern hematopoietic reference dataset, and two public erythroid cell reference datasets, to carefully characterize the cell types in the iPSC erythroid differentiation system. The cell map and cell differentiation trajectory of the system were mapped for the first time, and the whole transcriptome dynamics from iPSCs to the erythroid lineage were delineated.

This study observed that iPSCs went through two key transformations during EB formation to hematopoiesis and erythropoiesis. The first transformation was EHT, in which iPSCs differentiated into the germ layer under the induction of BMP4 and bFGF to form endothelial cells in the lateral plate mesoderm. Endothelial cells had strong cell-cell contact between themselves and other cells through the ESAM in the EHT. Under the induction of VEGF, HE was formed with the high expression of *FLI*, *LMO2*, and the cell adhesion molecules, and HE is converted to HSPCs with the up-regulated expression of *GATA2* under the influence of *RUNX1*, and the down-regulated expression of cell adhesion molecules. Reports showed that cell adhesion plays an essential regulatory function in mammalian embryonic development. Furthermore, cell-cell interactions were extremely indispensable in the information and the function of erythroblast islands, the niche where macrophages and immature erythrocytes were gathered and contacted to help erythrocytes maturation and enucleation during erythropoiesis [83–85].

The second transformation was HSPCs differentiation into erythroid cells. The HSPCs produced in the system were similar to CD34^+^, CD45^+^, and CD44^+^ YSMP [86], and they tend to differentiate into erythroid cells under the induction of EPO. YSMP enters the erythroid differentiation pathway driven by GATA1. During this period, the expression of estrogen target genes was up-regulated at the beginning and then decreased. A recent study revealed that estrogen could promote self-renewal of HSCs, expand splenic HSCs, and promote erythropoiesis during pregnancy [87]. A small amount of immune CD74^+^ erythroid cells are produced. Immune erythrocytes and their surface markers have been discovered and identified in recent years [88, 89] . This study found for the first time that the *in vitro* red blood cell regeneration system can also produce such cells, which provides a new direction for the clinical availability of RBCs regenerated *in vitro*. Also, we found that TFDP1 may be related to the restriction of cell number during the maturation of erythroid cells [65]. It was identified that *FOXO3* play a vital role in the denucleation of erythroid cells, and the high expression of *UPK1A-AS1* and *TRAPPC3L* in ortho-E may also regulate RBC denucleation-related genes.

Our results show that iPSCs go through two essential transformations to form erythroid cells in our system, accompanied by a complex TFs network and the dynamic expression of cell adhesion molecules and estrogen-responsive genes. It suggests that we can add cell adhesion activators before entering the first transformation, but add cell adhesion inhibitors during the transformation and inhibit endothelial TFs (e.g., SOX7) from increasing the production of HSPC in the system. Subsequently, we could activate erythroid TFs (e.g., GATA1) and add estradiol during the second transformation to increase the production of erythroid cells. This study provided a powerful theoretical guide for optimizing our iPSC-derived RBC regeneration system.

In summary, using the entire iPSC-derived RBC differentiation system in which both EB cells and SCs are included, we systematically characterized the molecular characteristics and differentiation trajectory of various cell types in the system. We provided essential factors driving iPSC differentiation into erythroid cells. However, technically it is not enough to improve the *in vitro* iPSC-derived RBC regeneration system at only one stage, and the RBC regeneration system needs to utilize different iPSC lines to prepare regenerative RBC of various blood types to suffice the needs of clinical blood transfusion under different blood types or different disease conditions in the future. Deciphering the RBC regeneration systems at different stages started from SCs in the EB culture system, we might identify more useful factors and pathways regulating adult globin expression and denucleation, which are the bottlenecks in this field. Moreover, we might characterize the real key regulators controlling RBC production process by comparing the *in vitro* iPSC-derived RBC regeneration system under physiological conditions (e.g., bone marrow). With the development of single-cell sequencing technology, we will be able to excavate more potential mechanisms regulating RBC development and maturation and obtain more mature regenerated functional RBCs *in vitro* for clinical applications in the future.

## Materials and methods

### iPSC and EB culture and erythropoiesis

iPSCs (sourced from Linzhao Cheng’s lab) were cultured in vitronectin (Invitrogen, A14700, Carlsbad, CA, USA)-coated culture dishes using essential 8 medium (Gibco, A1517001, Carlsbad, CA, USA). When the cells reached 70% to 80% confluence (usually around 3 to 4 days), they were digested to SCs using 0.5 mM EDTA (Invitrogen, 15575020, Carlsbad, CA, USA) and passaged according to a ratio of 1 to 4.

EB culture was performed on 0-11 d, as previously described [8]. From day 11 to 14 of the EB culture, TPO was replaced by 3 U/mL erythropoietin (EPO) (PeproTech,100-64, Cranbury, NJ, USA). On day 14, SCs were collected using a 70 μm cell strainer. Subsequently, the SC solution was centrifuged at 300 *g* for 5 minutes to remove the supernatant and SFM containing 50 μg/ml stem cell factor (PeproTech, 300-07, Cranbury, NJ, USA), 100 μg/ml IL-3 (PeproTech, 200-03, Cranbury, NJ, USA), 2 U/ml EPO, and 1% penicillin/streptomycin (P/S) (Life Technologies, 15140122, Carlsbad, CA, USA) was used to resuspend the SC precipitates. The cell concentration was adjusted to 5 × 10^5^/ml and cultured in a 6-well plate at a total volume of 2 ml per well. All cells were cultured for 6 days at 37°C under 5% CO^2^, and the medium was changed every three days. On day 20, the medium was replaced with SFM containing 2 U/ml EPO and 1% P/S, and the cell concentration was adjusted appropriately. The number of cells in each well did not exceed 2 × 10^6^, and the total volume of the medium per well was 2 ml. The cells were cultured at 37°C under 5% CO_2_, and the medium was changed every three days.

### Colony formation unit assay

Incubation of day 14 SCs was performed using MethoCult™ H4434 classic methylcellulose medium for human cells (StemCell, 04434, Vancouver, B.C., Canada). Each well of a 12-well plate contained around 10,000 cells and 0.6 ml of the medium. Other operations were performed as described in the instructions. Cells were incubated for 12-14 days, and the clones were observed under the microscope.

### Flow cytometry analysis

First, approximately 1 × 10^5^ cells were collected from the culture, centrifuged for five minutes at 300 *g* to remove the supernatant, and washed twice with 2 ml 1× Dulbecco’s phosphate-buffered saline (DPBS) (Gibco, 14190136, Carlsbad, CA, USA) solution containing 2% fetal bovine serum (Gibco, 16000044, Carlsbad, CA, USA) and 2 mM EDTA. After centrifugation the cells were resuspended in 50 μl DPBS buffer, and the antibodies (0.1 μl anti-CD71 and 0.5 μl anti-CD235a) were added, followed by incubation at 4°C for 10 min in the dark. The cells were washed twice by adding 2 ml of 1× DPBS buffer after incubation and resuspension in 200 μl of DPBS buffer to prepare a cell suspension for the assay. The BD FACSAria II instrument was used for flow cytometry analysis, and the data analysis was performed with Flowjo software (v.7.6, Three Star). The antibodies (Miltenyi Biotec, Bergisch Gladbach, Germany) used in this study included anti-human CD34-FITC (130-098-142), anti-human CD45-PE (170-081-061), anti-human CD71-PE (130-099-219), and anti-human CD235a-APC (130-118-493).

### Wright-Giemsa staining

Approximately 1 × 10^5^ cells were taken, and the medium was removed by centrifugation at 300 *g* for 5 minutes; the cells were resuspended in 20-μl DPBS buffer. The cells were placed onto one of two clean slides, and the second slide was used to push the first to 30°-45° to spread the cells evenly. The pushed film was placed in a 37°C incubator for one to three hours. Then, 200 μL of Wright-Giemsa staining A solution (BASO, BA-4017, Zhuhai Guangdong, China) was placed onto the cell-containing slide, and the slide was placed at room temperature for one minute. It was immediately covered with 400 μL of B solution. After the slide was incubated at room temperature for 8 minutes, the slides were gently rinsed with fluid distilled water along the area around the slide without cells. The slides were rinsed for an additional 1 minute until the water ran clear and transparent. Then, the slides were placed at room temperature until dry, and a microscope was used to observe and record cell morphology.

### Total RNA extraction and qPCR

Total RNA was extracted from the iPSC line BC1, day four EBs, day 14 SCs, and day 20 erythroid cells, and the total RNA was reverse transcribed into cDNA using the PrimeScript RT Reagent Kit with gDNA Eraser (TaKaRa, RR047A, Kusatsu, Shiga, Japan). The diluted cDNA was used as a template for qPCR. The instrument used for qPCR was the Bio-Red CFX96 Real-Time PCR detection system. All primers used in qPCR are listed in Table S2.

### scRNA-seq data analysis and bulk RNA-seq data analysis

Day 14 EBs were dissociated with Accutase solution (Sigma, A6964, St. Louis, Missouri, USA). Single-cell suspensions at 800 cells/µL were subjected to chromium 10× genomics library construction. Raw gene expression matrices generated by CellRanger (v.3.0.0) were imported into Seurat (v.3.2.2) [46]. After removing low quality cells and genes (percentage of mitochondria reads ≥ 25%, unique genes ≤ 200 in a cell, and genes expressed less than 5 cells), the left cells were annotated by SingleR and classified according to annotation results [48]. TFs regulatory network was constructed by SCENIC (v.1.1.2) [63], and cell contact patterns were constructed by CellChat (v.0.0.2) [60]. Particular clusters were imported into Monocle2 [66] for pseudo-time ordering. For bulk RNA-seq data, FASTQC and Trimmomatic were used to remove adaptor sequences and low-quality reads. Then, clean data were mapped to the reference genome (GRCh38) by HISAT2 to generate a gene expression matrix, which was then imported into R Package DEGseq to identify DEGs. Heatmap analysis and PCA were performed by pheatmap and PRINCOMP, respectively. We used clusterProfiler (v.3.18.0) to perform functional annotation analysis [59]. All data analyses were completed in R (v4.0.0) environment. The list of human TFs was downloaded from http://humantfs.ccbr.utoronto.ca/download.php. The Circular network was visualized by GeneMANIA [90].

## Data availability

The single-cell RNA-seq of D14 EBCs and SCs, and the bulk RNA-seq of BC1 and D14 SCs is available in the Genome Sequence Archive (GSA) http://bigd.big.ac.cn/gsa/s/ZVZPnz5q

## CRediT author statement

**Xinzi Juan:** Investigation, Data curation, Formal analysis, Validation, Visualization, Writing - original draft, Writing - review & editing. **Wei Zhang:** Investigation, Data curation, Formal analysis, Validation, Visualization, Writing - original draft, Writing - review & editing. **Shangjin Gong:** Formal analysis. **Junwei Zhu:** Validation. **Yanming Li:** Project administration. **Zhaojun Zhang:** Conceptualization, Resources, Writing - original draft, Writing - review & editing, Supervision. **Xiangdong Fang:** Conceptualization, Resources, Writing - original draft, Writing - review & editing, Supervision.

## Acknowledgments

This research was supported by the Strategic Priority Research Program of the Chinese Academy of Sciences (Grant No. XDA16010602), the National Key Research and Development Program of China (Grant Nos. 2016YFC0901700, 2017YFC0907400, and 2018YFC0910700), and the National Natural Science Foundation of China (Grant Nos. 81670109, 81870097, 81700097, 81700116, and 82070114). We were grateful to Linzhao Cheng for agreeing to use iPSC line BC1 and Qianfei Wang for providing the iPSCs. We thank the editor Suzanne N in enago editing service who provided assistance for the manuscript.

## Supplementary material

**Figure S1 Comparison of two batches**

**A**. Pie chart of the cell number percentage of the two batches. **B**. Correlation analysis of the two batches by the Spearman correlation coefficient of all detected genes.

**Figure S2 Cell score heatmap of 32 cell types annotated by SingleR and the distribution of each cell type in the two sample batches.** Batch 1 is represented in orange, and batch 2 is represented in blue.

**Figure S3 The location of each cell type annotated by SingleR in the UMAP plot.**

**Figure S4 Each cell classification counts in batches and correlation analysis**

**A**. The number of cells in each cell classification after cell-type classification (left) and the batch source of each cell classification (right). **B**. Heatmap plot of Spearman correlation coefficient between cell classifications.

**Figure S5 Cell-cell contact involving bulk RNA-seq data analysis**

**A** and **B**. The cell-cell contact patterns of iPSCs-derived erythrocyte regeneration system, cell patterns_outgoing (**A)**, Cell patterns_incoming (**B)**. **C**. Heatmap plot of hematopoietic lineage-related gene expression patterns for four bulk RNA-seq datasets. Blue indicates low gene expression and red indicates high gene expression. **D**. PCA of four bulk RNA-seq datasets. The red arrow indicates the PCA position of the four bulk RNA-seq datasets. **E**. Gene function annotation analysis for low expression genes in SCs.

**Figure S6 GO annotation of target genes in regulons**

The GO enrichment results and heatmap plot displayed the relationship between biological processes and the target genes of TFs in Figure 4C. NANOG and POU5F1 that are activated in starting cells (**A** and **B**), SOX10, ONECUT2, and SIX1 that are activated in ectoderm-derived cells (**C** and **D**), FOXA3, CREB3, and TEAD1 that are activated in intermediated mesoderm-derived cells (**E** and **F**), HOXC6 and PRDM6 that performed functions in paraxial mesoderm-derived cells (**G** and **H**), MECOM, ETS1, ELK3, ERG, and SOX18 that performed functions in lateral mesoderm-derived cells (**I** and **J**), GATA1, TAL1, RUNX1, CEBPD, MYB, and ELF1 that are activated in hematopoietic cells (**K** and **L**). In contrast, CEBPD and MYB were specifically activated in myeloid cells (**M** and **N**) and GATA1 and TAL1 were specifically activated in the erythroid cells (**O** and **P**).

**Figure S7 Distribution of regulons activity in TFs UMAP reduced dimension** Schematic diagram of the regions of 21 cell classifications specificity activation regulons on the UMAP map. Regulon activated cells were colored red, and regulon non-activated cells were colored gray.

**Figure S8 RSS (regulon specificity score) pattern of each hematopoietic cell type** The top five regulons in each hematopoietic cell type were highlighted in red and labeled on the plot based on the RSS. The content in the label brackets is the number of target genes in the regulons. The regulons specificity score is shown on the y-axis.

**Figure S9 Regulation of iPSCs differentiation direction**

**A**. The arrangement of each classification cells on pseudo-time trajectory. **B**. *POU5F1*, *RUNX1*, and *SOX7* expression level on total cell pseudo-time trajectory. **C**. KEGG analysis of marker genes in each state cell. **D**. Volcanic plot of DEGs from gene differential expression analysis between state two and three initial cells (*P* < 0.05, |log2 FC| < 0.5). **E**. Identification of top20 TFs regulating high expression genes in state two initial cells. **F**. Network of TFs from E by GeneMANIA.

**Figure S10 Dynamic expression of cell adhesion molecules during EHT**

**A**. The arrangement of embryonic stem cells, iPS cells, endothelial cells, and CD34^+^ HSPCs on pseudo-time trajectory. **B**. The arrangement of state cells on pseudo-time trajectory. **C**. The expression of trend changes between states of each cluster gene from Figure 5C. **D**. qPCR analysis of cell adhesion molecules showing fluctuation throughout EHT.

**Figure S11 Hematopoietic cells differentiation trajectory**

**A**. UMAP plot of hematopoietic cells distribution. **B**. Cell proportion pie chart of hematopoietic cells. **C**. The arrangement of each type of cells on hematopoietic pseudo-time trajectory. **D**. *GATA1*and *CEBPD* expression level on hematopoietic cells pseudo-time trajectory. **E**. The expression of trend changes between states of each cluster gene from Figure 7E.

**Figure S12 Dynamic expression of estrogen target genes during erythroid differentiation**

**A**. The arrangement of BFU-E, CFU-E, erythrocytes, ortho-E cells on erythroid pseudo-time trajectory. **B**. Pseudo-time analysis of genes in the estrogen signaling pathway KEGG term from Figure 6C. **C**. GO analysis of cluster 2 genes from Figure 8C. **D**. The expression of trend changes between states of each cluster gene from Figure 8C. **E**. qPCR analysis of the effects of estradiol during erythroid differentiation. **F**. Volcanic plot of DEGs from gene differential expression analysis between state one and three cells (*P* < 0.05, |log2 FC| < 0.5). **G**. Volcanic plot of DEGs from gene differential expression analysis between ortho-E and other erythroid cells (P < 0.05, |log2 FC| < 0.5). **H**. KEGG analysis of DEGs from **G**.

**Table S1** Sample quality evaluation indicators and results

**Table S2** qPCR primer for globin, cell adhesion and estradiol related genes

